# Mechano-signaling of prostate tumor initiating cells facilitates their tropism to stiff metastatic niche

**DOI:** 10.1101/2023.08.28.553410

**Authors:** Lanpeng Chen, Gangyin Zhao, Marta De Menna, Stefano Coppola, Nick Landman, Sebastiaan Schieven, Arwin Groenewoud, George N. Thalmann, Thomas Schmidt, Jelle de Vries, Marianna Kruithof-de Julio, Ewa B. Snaar-Jagalska

## Abstract

Analysis of clinical datasets indicate that cancer stem-like cells/tumor-initiating cells (CSCs/TICs) derived from prostate cancer (PCa) patients display an elevated expression of genes for cell-matrix interactions, cell adhesion proteins and of the putative mechanotransducer TAZ. Here we combined measurements on the cellular mechano-responses to matrix stiffness, including cell-generated forces, zebrafish and PDX-derived organoid models, to show that mechanotransduction serves as a key determinant for PCa CSC maintenance during metastatic onset. The β1-integrin-ILK-CDC42-N-Wasp dependent cytoskeletal tension and TAZ nucleus-translocation mediate this mechano-signaling axis. As a result, expression of the stemness genes NANOG and OCT4 are induced, leading to metastatic tumor initiation. It is further demonstrated that pharmaceutical perturbation of this mechano-signaling using a novel YAP/TAZ inhibitor K975 constrains PCa metastasis in zebrafish, and development of PDX-derived organoids. Our data highlights the essential role of mechanotransduction in PCa aggressiveness, thereby underlying this pathway as a therapeutic target for future studies.

---

Incurable bone metastasis is the main cause of death in prostate cancer (PCa) (1). It is suggested to be initiated by cancer stem-like cells (CSCs) or tumor-initiating cells (TICs) (2–5), a small cell subpopulation characterized by enhanced mobility, self-renewal capacity, tumorigenicity and chemo-resistance. Therapeutic strategy against this cell population, however, is still missing. Therefore, further to understand molecular and cellular mechanisms of CSC regulation to identify potential molecular compounds targeting this cell population is of great interest. Previous studies show that CSCs derived from PCa patients displayed a significantly enhanced expression of gene signatures for various focal adhesion proteins (6). Hence, the cell adhesion molecules CD44 and integrin α2β1 have been used as PCa CSC markers (7–10). Both proteins are functionally involved in PCa progression, through the strengthening of focal adhesions, altering cytoskeleton dynamics, and built-up of cytoskeletal tension, implicated involvement of their cell mechanics in the initiation of metastasis, mainly into stiff bone (2,6,11).

Mechanotransduction is a process in which external mechanical cues from the microenvironment are transduced into biochemical signals inside the cells through focal adhesion and actin cytoskeleton remodeling (12,13), modulating gene expression and metabolisms (14,15). A key of this cellular mechano-response is the transcriptional regulator YAP/TAZ (16–20). Once activated by mechanical cues, YAP/TAZ translocates into the nucleus and thereby drives gene expression through binding to multiple transcriptional factors that e.g. control cell survival, proliferation, cell fate, and metabolic wiring (15,17,21). Importantly, YAP/TAZ plays an essential role in stemness regulation during embryonic development (22–27). Hence, transient activation of YAP/TAZ by mechanical stimulation can reprogram differentiated cells into their corresponding stem or progenitor status (26). In cancer, active YAP/TAZ has been reported to promote cancer stemness(28) in breast cancer, osteosarcoma, and (29) glioblastoma (20,30–32), suggesting YAP/TAZ could serve as a potential target against CSCs.

In order to assess the role of mechano-responses in the regulation of PCa cancer stemness and metastasis, and to identify potential molecular compounds specific for CSCs targeting, we measured mechanical responses of PCa cells to matrix stiffness, revealing that the metastatic potential of PCa CSCs relies on their mechanosensing of ECM stiffening and mechanotransduction ability. This cell mechanic is mediated by integrin β1 and integrin-linked kinase (ILK) that lead to significant cytoskeleton remodeling through CDC-42-N-Wasp axis. This further increases cytoskeletal tension that results in nuclear translocation of TAZ (29,33,34). The latter driving expression of pluripotency genes, a prerequisite to metastatic tumor initiation. Suppression of this signaling cascade using a novel YAP/TZA inhibitor, K975 (35), significantly attenuated metastasis in zebrafish xenografts and blocked near-patient organoid growth. Overall, our study underscores that mechanotransduction of CSC endows their tropism, repopulation and metastatic initiation in stiff niche to served as a key determinant of CSC regulation in metastasis, and suggesting the pathway as future therapeutic target.

## Results

### PCa CSCs have enhanced capacity of mechanosensing and mechanotransduction

To address if mechanotransduction could mediate PCa metastasis, we first compared the expression of the mechano-transducer YAP and TAZ in patients’ metastases with that in primary tumors in 2 different clinical datasets (Taylor Prostate (181 primary tissues and 37 metastasis tissue) and Grasso Prostate (59 primary tissues and 35 metastasis tissues) (36,37)). In both datasets, TAZ were significantly upregulated in metastasis compared to that in primary tumor (Fig. 1a). In addition, we compared YAP and TAZ expression in patients’ derived CSCs versus non-CSCs in a public accessible dataset (2), showing that TAZ but not YAP was increasingly expressed in the CD133 ^+^CD44^+^α2β1^+^ CSCs compared to their CD133 ^-^CD44^-^α2β1^-^ non-CSC counterparts (Fig. 1b). Overall, this transcriptomic analysis suggests a close correlation between TAZ expression, CSCs and metastasis.

**Fig. 1.**
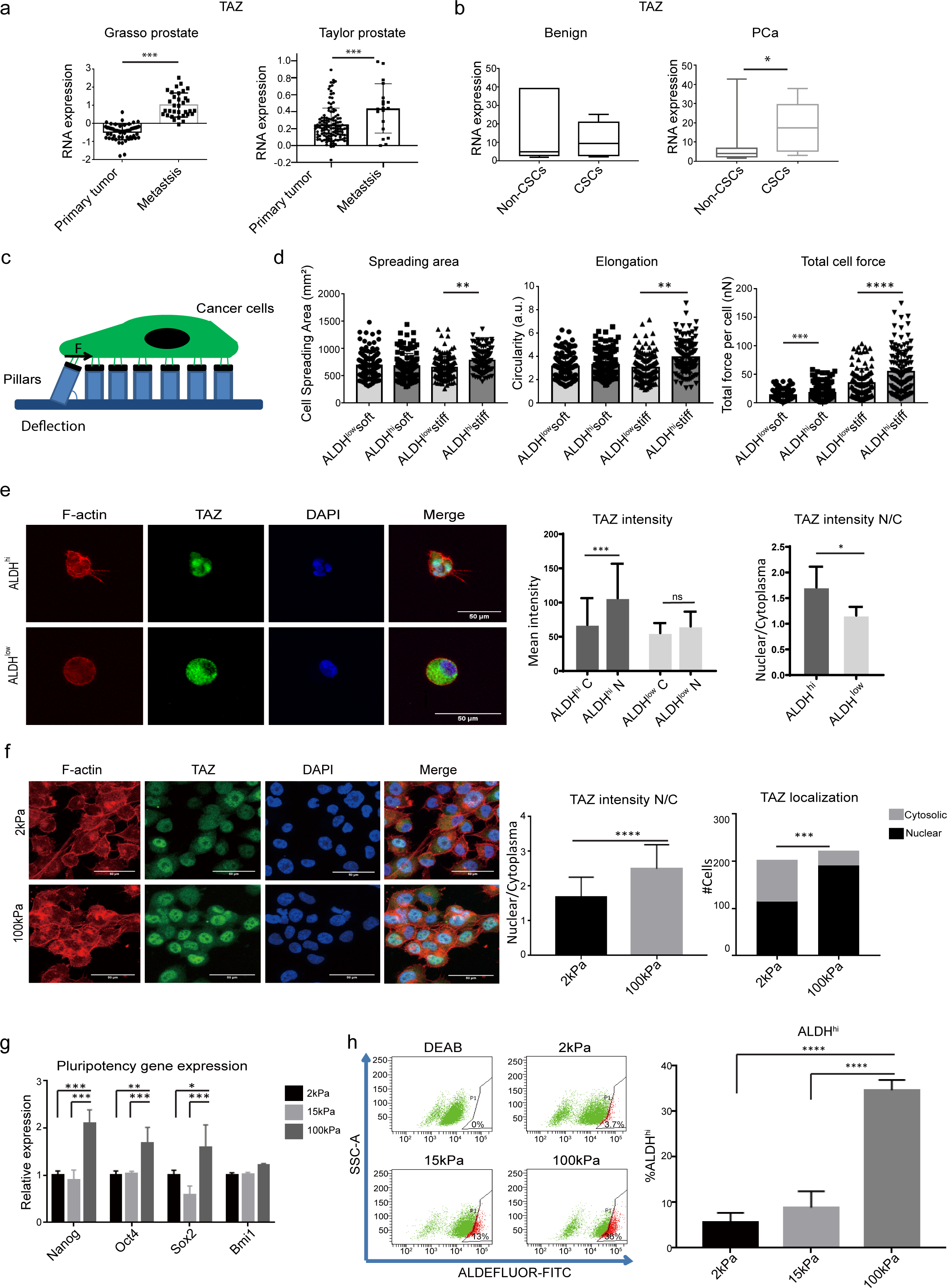
PCa CSC-like phenotype correlates with enhanced focal adhesion, cell contractility and TAZ nuclear translocation. (a) TAZ expression was compared between PCa metastases and the primary site. Clinical datasets Taylor Prostate (GSE21032) and Grasso Prostate (GSE35988) were used for the analysis (data accessed from www.oncomine.org). (b) Comparison of TAZ expression between patients’ derived CSCs and non-CSCs in a published dataset (E-MEXP-993). (c-d) ALDH ^low^ and ALDH^hi^ subpopulations were seeded on elastic micro-pillar arrays with low (28kPa) and high (142kPa) stiffness for 24 hours. Cell spreading area, Elongation and Contractile force were measured. 120 cells from 3 independent experiments were analyzed for each group. (e) ALDH ^low^ and ALDH^hi^ subpopulations were seeded in MoT 3D culture. Immunofluorescence against TAZ was conducted. TAZ intensity in nucleus and in cytoplasm was calculated. (f) PC-3M-Pro4 cells were seeded on elastic cell culture substrates with stiffness of 2kPa and 100kPa for 48 hours. Immunofluorescence against TAZ was performed. TAZ signal intensity in cytoplasm and nucleus was analyzed using an Image-J program. Group size = 30. (g-h) PC-3M-Pro4 were seeded on 2kPa, 15kPa and 100kPa for 96 hours. Expression of pluripotency genes (g) and size of ALDH ^hi^ subpopulation (h) and were measured. Experiments were independently repeated 3 times. Scare bar = 50um. Group size = 30. *p<0.05, **p<0.01, ***p<0.001, ****p<0,0001. Arrow bars are s.e.m.

To experimentally address if PCa CSCs have higher mechanotransduction capacity, CSC-like cells were sorted from an osteotropic PCa cell line PC-3M-Pro4 using ALDFLUOR assay, a well-established method for selection of CSC-like cells based on their Aldehydedehydrogenase (ALDH) activities (38). Directly after sorting, the ALDH ^hi^ CSC-like cells and their ALDH^low^ counterparts were respectively seeded on fibronectin-coated elastic micropillar substrate with a low (28kPa) or high (142kPa) stiffness (39). Cell spreading area, elongation and contractile force generated by the cell-ECM interaction were measured (Fig. 1c). Compared to the ALDH ^low^ population, the ALDH ^hi^ cells showed elevated spreading and elongation on the rigid substrate associated with a significantly enhanced contractile force generation (Fig. 1d). Next, the ALDH ^hi^ and ALDH^low^ cells were seeded for Matrigel on top (MoT) 3D culture, an *in vitro* 3D assay to mimic cancer cell metastatic colonization (40). We compared TAZ nuclear translocation between the two subpopulations: ALDH ^hi^ cells had significantly elevated TAZ nuclear translocation accompanied by enhanced formation of Filopodia-protrusions (FLPs) in comparison to the ALDH ^low^ (Fig. 1e, Fig. S1a). Taken together, these results confirmed that PCa CSCs display enhanced mechanosensing and mechanotransduction capacity leading to TAZ activation.

The increased mechano-sensing and -transduction capacities in the ALDH^hi^ cells prompt us to question if ECM stiffness can impact size of^hi^ tchelels AinLtDheHbulk population. We therefore seeded bulk PC-3M-Pro4 on elastic substrates with defined stiffness: 2kPa, 15kPa (data not shown) and 100kPa. Activation of the mechanical signaling was determined by measuring TAZ nuclear translocation using immunofluorescence. As expected, the ratio of nuclear to cytosolic TAZ intensity was significantly elevated in the cells on 100kPa compared to the 2kPa (p<0.001) (Fig. 1f), indicating that PCa cells can respond to the increased stiffness and induce TAZ nuclear translocation. Following the increase of ECM stiffness, expression of the stemness genes *NANOG*, *OCT4* and *SOX2* w a s significantly increased (Fig. 1g), and the size of the ALDH^hi^ subpopulation was elevated to 30% on 100 kPa from 5% on 2kPa and 9% on 15kPa (Fig. 1h).

### Inhibition of mechanotransduction by targeting of Integrin β1-ILK-TAZ axis suppresses CSCs *in vitro*

Having shown the increase of ECM stiffness could enhance TAZ nuclear translocation and stem-marker expression, we next questioned how the cells sensed the stiffness of the ECM and transmit the signal into the cells. Given that the focal-adhesion transducer and cytoskeleton regulator Integrin β1 (encoded by *ITGB1*) (41) and its interactor Integrin-Linked Kinase (*ILK*) were significantly upregulated in the ALDH ^hi^ subpopulation (Fig. S1b-S1c), we tested if this integrin signaling could mediate mechanotransduction and thus controlled the stem-like phenotype of the cells. To this end, RNA interference approach was applied to inhibit *ITGB1* and *ILK* expression. Both knockdowns strongly inhibited Paxillin phosphorylation (a marker of focal adhesion) and formation of FLPs in the ALDH^hi^ cells, underlining the involvement of integrin signaling in focal adhesion formation and actin polymerization (Fig. S1d). We subsequently assessed the effects of *ITGB1* kd and *ILK* kd on the CSC markers (Fig. 2a-2b). Both knockdowns significantly suppressed the expression of pluripotency genes *NANOG*, *OCT4* and *BMI1* by more than 70% (p<0.001) (Fig. 2a), concomitant with a significant reduction of the ALDH^hi^ subpopulation compared to the scramble shRNA (SCR) control (Fig. 2b). Apart from the suppression of the stemness markers, *in vitro* tumorigenicity of the cancer cells was also impaired by the knockdowns as indicated by the clonogenicity and tumor spheroid assays in both PC-3M-Pro4 and C4-2B cell lines (Fig. 2c, S2a-S2c), demonstrating integrin β1 and ILK were functionally involved in regulation of the CSC-like phenotype.

**Fig. 2.**
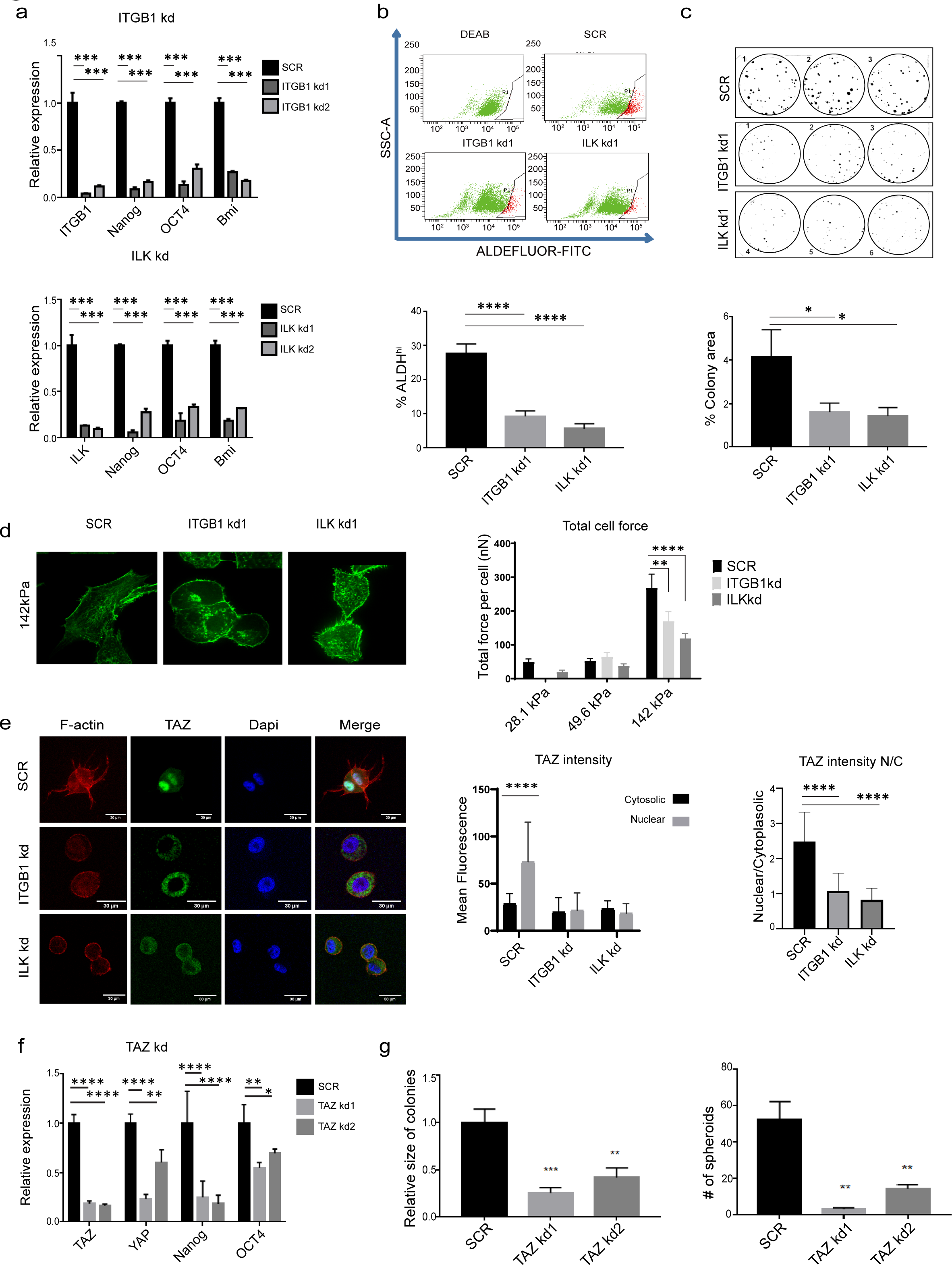
Knockdowns of integrin β, ILK and TAZ inhibit PCa CSC-like phenotype. (a-c) Effects of integrin β1 (ITGB1) and ILK knockdowns on the CSC-like phenotype of PC-3M-Pro4. The expression of pluripotency genes (a), size of ALDH^hi^ fraction (b) and formation of colonies (c) were tested. (d) Knockdowns of integrin β1 and ILK suppress cell contractility on high stiffness. (e) Knockdowns of integrin β1 and ILK suppress TAZ nuclear translocation. (f) Effects of TAZ knockdowns on pluripotent gene expression (g), clonogenicity and tumor spheroid formation of PC-3M-Pro4 *in vitro.* E a c h experiment was independently repeated 3 times. *p<0.05, **p<0.01, ***p<0.001, ****p<0,0001. Arrow bars are s.e.m.

Next, we measured contractile force generated by the cancer cells bearing SCR control*, ITGB1* kd and *ILK* kd by seeding the cells on micro-pillars array with defined stiffness of 28.1 kPa, 49.6kPa and 142kPa. At 24 hours after seeding, cell contractile force was calculated based on the pillars deflection. Following the increase of stiffness, contractile force generated by the SCR cells was significantly elevated (from 50nN on 28.1 and 49.6 kPa to 280nN on 142kPa (Fig. 2d)), while the knockdowns of *ITGB1* and *ILK* significantly reduced contractile force of the cells on 142kPa, suggesting integrin β1 and ILK are key regulators for cell contractility (Fig. 2d). Consistent with this, when the cells were cultured on matrigel, the formation of FLPs (Fig 2e, S2d, S2e) and nuclear translocation of TAZ (and YAP) (Fig 2e, S3a, S3b) were significantly attenuated by the knockdowns of Integrin β1 and ILK, accompanied by suppression of TAZ downstream genes: AMTOL, CTGF and CYR61 (Fig. S3c). This suggests that Integrin β1 and ILK1 may play a bridging role in regulating the nuclear localization of TAZ and cytoskeletal remodeling.

We next tested if the integrin β1-ILK axis modulates the CSC-like phenotype through TAZ activation. It showed that interference of TAZ (and YAP) by shRNA abrogated expression of the pluripotent genes *NANOG* and *OCT4* in both PC-3M-Pro4 and/or C4-2B cell lines (Fig. 2f and Fig.S3d), and attenuated their capacity of colony formation and tumor spheroid in 2D and 3D culture respectively (Fig. 2g), confirming that the mechano-signaling governed by the integrin β1-ILK-TAZ axis is required for maintenance of the CSC-like phenotype in PCa.

### Disruption of the integrin β1-ILK-cytoskeleton-TAZ axis inhibits metastatic onset *in vivo*

Having shown roles of the integrin β1-ILK-TAZ axis in regulation of the CSC-like phenotype *in vitro*, we assessed if this signaling axis regulates PCa metastasis initiation in a zebrafish (ZF) xenograft model (42–44). In order to monitor actin cytoskeleton remodeling *in vivo,* PC-3M-Pro4 and PC-3 cells were transfected with lifeact-mcherry, a small protein peptide specifically labeling actin filaments. Intravenous transplantation of the PCa cells into ZF larva at 2 days post fertilization (2dpf) led to cancer cell dissemination following circulation. At 1 days post injection (1dpi), the disseminated cells adhered to caudal vein (labeled with eGFP) and extravasated into caudal hematopoietic tissue (CHT), where the cells started to grow and invaded the neighboring tissues in the following 5 days (Fig. 3a, Fig. S4a). This process was accompanied intensive cytoskeleton remodeling, characterized by the formation of fillopodia-like and invadopodia-like protrusions and focal adhesion plague (marked by localization of Paxillin at cell edge) (Fig. S4b, S4c). Additionally, TAZ nuclear translocation occurred (Fig. S4d), indicating a mechanistic link between actin dynamics and TAZ activation in the metastatic colonization of the engrafted cells.

**Fig. 3.**
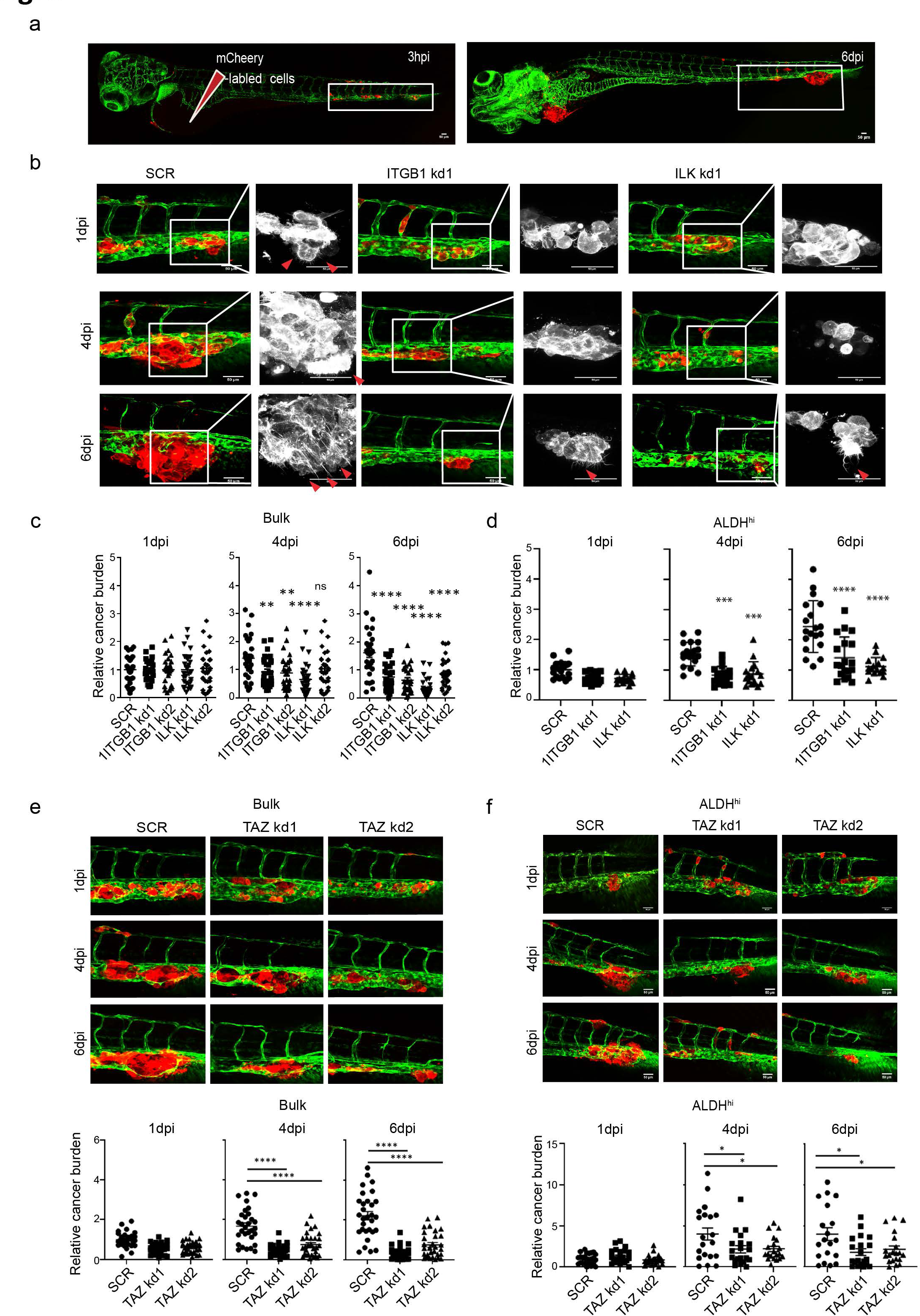
Knockdowns of integrin β1, ILK and TAZ suppress metastatic onset of PCa cells in the zebrafish xenografts. (a) Scheme of intravenous transplantation of PCa cells into ZF. PCa cells were stably expressed with Lifeact-mCherry and injected into duct of cuvier (DoC) at 2 days post fertilization (2dpf). Perivascular metastases at CHT area (white box) was developed at 6 days post injection (dpi) (red, cancer cells; green ZF vessel). (b) PC-3M-Pro4-lifeact-mCherry-SCR,-ITGB1 kd and -ILK kd were respectively injected into ZF. Images were acquired at 1, 2, 4 and 6 dpi. Red arrow, Actin filaments. Scale bar=50um. (c) Growth of the PC-3M-Pro4 cells bearing SCR, ITGB1kd and ILKkd at the metastatic site were analyzed. Group size=30. Experiments were independently repeated 2 times. (d) ALDH^hi^ subpopulation was isolated from PC-3M-Pro4-Lifeact-mCherry-SCR, -ITGB1 kd1 or -ILK kd1 and injected into ZF. Growth of the cancer cells at metastatic site were analyzed. Group size= 20. (e-f) Growth of the bulk (e) and ALDH ^hi^ (f) PC-3M-Pro4 cells bearing SCR, TAZ kd1 and TAZ kd2 at metastatic site were analyzed. Group size= 30. *p<0.05, **p<0.01, ***p<0.001, ****p<0,0001. Arrow bars are s.e.m.

Next, we elucidated if this is also valid for the ALDH ^hi^ subpopulations. Indeed, after transplantation, the ALDH ^hi^ cells displayed enhanced cytoskeleton remodeling, extravasation and metastatic growth at CHT, conforming that ALDH ^hi^ cells bear increased metastatic potential in ZF (Fig. S4e), in line with previous findings (45,46).

In order to test if the mechano-signaling is required for the metastatic colonization, we engrafted the PCa cells containing SCR control, ITGB1 kd and ILK kd into ZF. Consequently, growth of the ITGB1 kd and ILK kd cells at the metastatic site was significantly inhibited by more than 50% accompanied by an attenuation of cytoskeleton remodeling in both PC-3M-Pro4 (Fig. 3b,c) and PC-3 (data not shown). To reveal if these knockdowns also impaired the metastatic potential of the ALDH^hi^ subpopulations, the ALDH ^hi^ cells were sorted from PC-3M-Pro4-SCR, -ITGB1 kd1 and -ILK kd1 cell lines and transplanted into ZF. As expected, metastatic growth of the ALDH ^hi^ subpopulation was significantly suppressed by the knockdowns (Fig. 3d).

Given that the reduction of metastatic tumor growth in the knockdown cells was associated with suppression of cytoskeleton remodeling, we questioned whether the Integrin-β1-ILK signaling axis governs metastasis by cytoskeleton regulation. To address this, we conducted an *in vivo* shRNA screening using the ZF model. In total 8 cytoskeletal remodeling regulatory genes were tested including CDC42, RAC1, LIM kinase 1 (LIMK1), Profilin1, Wasp, N-Wasp, Cortactin and DIAPH3 (mDIA2) (Fig. S5a). Six out of eight knockdowns significantly suppressed metastatic growth of the cells (Fig. S5b). Notably, the knockdowns of CDC42 and its downstream proteins N-Wasp and Cortactin showed the strongest inhibitory effect at 6dpi (inhibition >50%, p<0.001) (Fig. S5a-S5b), concomitant with reduced cytoskeleton remodeling (Fig. S5c). In order to further ascertain that the disruption of cytoskeleton suppressed metastasis by blocking initiation of tumor growth, we transduced PC-3M-Pro4 bearing CDC42 and N-WASP knockdowns with Fluorescent ubiquitination-based cell cycle indicator (Fucci), and injected those cells into zebrafish. At 1 dpi, all groups had equal percentage of cells at G0/G1 (red) and G2/M (green) phase. At 4 dpi, a significant increase in the percentage of cells at the G2/M phase was observed in the SCR control (Fig. S5d), indicating the initiation of tumor growth at the metastatic site. However, this initiation was inhibited by knockdowns of CDC42 and N-Wasp, as evidenced by a low percentage of cells in the G2/M phase, suggesting that these cytoskeletal proteins are essential for metastatic tumor growth (Fig. S5d). In addition, *in vitro* approaches showed the knockdown of CDC42 and N-Wasp attenuated the size of ALDH^hi^ cell subpopulation, lowered the expression of stemness genes *NANOG* and *OCT4*, and suppressed tumorspheroid formation and clonogenicity (Fig. S6a-S6c), simultaneously with reduction of TAZ nuclear translocation and TAZ downstream gene expression (Fig. S6d). Altogether, this data indicates that the cytoskeleton signaling pathway is required for the TAZ-dependent CSC maintenance.

Lastly, we determined if the disruption of the integrin-cytoskeleton signaling axis suppressed metastasis by the inhibition of TAZ. We showed that the TAZ knockdowns significantly suppressed metastatic growth of PC-3M-Pro4 at 4 and 6 dpi (Fig. 3e). In addition, the ALDH^hi^ subpopulation sorted from the TAZ knockdown cells had significantly reduced metastatic capacity in ZF, indicating that the TAZ activation, putatively driven by the integrin-dependent cytoskeleton remodeling, governs the aggressiveness of the ALDH ^hi^ cells (Fig. 3f).

### Overexpression of integrin β1, ILK and TAZ-S89A results in enhanced stem-like phenotype in PCa cells

In order to further confirm the determinant role of the integrin-TAZ signaling axis in controlling the stem-like phenotype and metastasis in PCa cells, we used lentiviral approaches to overexpress ITGB1, ILK and constitutively active TAZ (TAZ-S89A) in PCa cell lines PC-3, PC-3M-Pro4 and C4-2B. Compared to the empty vector (Mock), overexpression of ITGB1, ILK and TAZ-S89A significantly elevated expression of the pluripotent genes NANOG, OCT4 and/or BMI-1 on both transcriptional and protein levels (Fig. 4a-b). Among all overexpression groups, the cells with TAZ-S89A overexpression showed the highest pluripotent gene elevation (Fig. 4a-b). In line with this observation, TAZ-S89A overexpression significantly increased the size of the ALDHhi fraction in all tested cell lines (Fig. 4c), including PC3 (from 6.81% to 23.6%), PC-3M-Pro4 (from 13.3% to 26.5%), and C4-2B (from 2.65% to 8.33%), indicating that TAZ is a key driver of CSC maintenance in PCa. We next test how the overexpression of ITGB1, ILK and TAZ impact metastatic capacity of the cancer cells in the zebrafish model. PC-3 and PC-3M-Pro4 bearing ITGB1, ILK or TAZ-S89A overexpression were I.V. injected into ZF. As expected, the overexpression significantly increased proliferation, extravasation and invasion of the engrafted cells (Fig. 4d). Notably, the cells overexpressed with TAZ-S89A displayed the highest increase (Fig. 4d, 4e), in line with its strong impacts on the stemness gene expression (Fig. 4a). Overall, this genetic study demonstrated that the integrin-TAZ signaling axis in PCa cells serves as a key determinant of cancer aggressiveness.

**Fig. 4.**
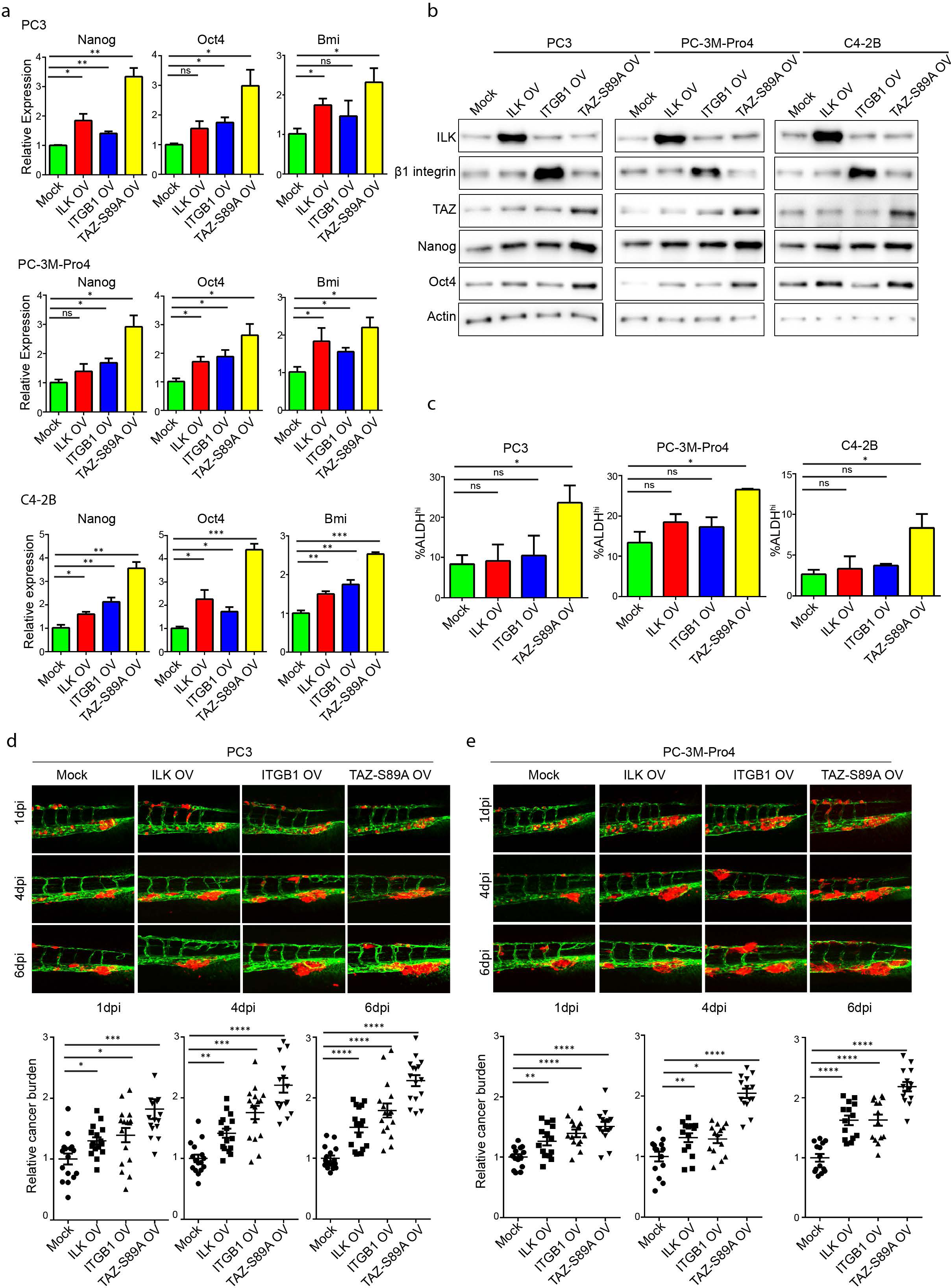
Overexpression of integrin β1, ILK and constitutively activated TAZ in PCa cells increases pluripotent gene expression and size of ALDH ^hi^ subpopulation in vitro, and metastasis in the zebrafish xenografts. (a-b) Overexpression of integrin β1, ILK and constitutively activated TAZ elevates pluripotent gene expression on transcriptional (a) and protein (b) levels in PCa cell lines. Each experiment was independently repeated 3 times. (c) Size of ALDH ^hi^ subsets in PCa cell lines overexpressed with empty vector (Mock), Integrin β1, ILK and constitutively activated TAZ. Each experiment was independently repeated 3 times. (d-e) Overexpression of integrin β1, ILK and constitutively activated TAZ increases metastasis of PC-3 (d) and PC-3M-Pro4 (e) in zebrafish xenografts. Group size = 30 in 2 independent experiments. *p<0.05, **p<0.01, ***p<0.001, ****p<0,0001. Arrow bars are s.e.m.

### TAZ inhibitor K975 suppresses pluripotent gene expression, size of ALDH^hi^ subpopulation and metastasis of PCa cells

The key role of integrin β1-ILK-TAZ signaling axis in regulation of the CSC-like phenotype elicited a hypothesis that targeting of this signaling can suppress PCa metastasis. To prove this, we tested serial small molecular compounds targeting the signaling axis including the integrin β1 inhibitor BTT3033, ILK inhibitor CPD22, YAP/TAZ inhibitor verteporfin, and K975, a novel TAZ inhibitor which covalently binds to an internal cysteine residue located in the palmitate-binding pocket of TEAD (47), and thereby blocks its activation by interacting with nuclear TAZ. Although all inhibitors exerted anti-cancer activities (data not shown) to distinct levels, K975 displayed the lowest cytotoxicity with a desired inhibitory effect (>50%) on TAZ downstream gene expression (Fig. 5a-c). We therefore decided to explore anti-cancer potential of this compound. When PC-3, PC-3M-Pro4 and C4-2B were treated for 48h with K975 at concentrations between 5 and 20 µM, expression of the pluripotent genes *NANOG*, *OCT4* and *BMI1* were significantly decreased (±50% inhibition at 10-20uM) (Fig. 5a-c), concomitant with a significant suppression of the ALDH^hi^ subset (Fig.5d). Next, anti-metastasis efficacy of K975 was tested in the ZF model. 0.5, 1 or 5 pmol K975 was I.V. injected into ZF at 4 hours post transplantation. Metastasis at CHT was monitored, showing that the K975 injection significantly inhibited metastatic tumor growth in a dose-dependent manner (±70% inhibition at 0.5 pmol and >90% inhibition at 5 pmol at 6 dpi) (Fig.5e and 5f). Overall, our experimental data demonstrate that the TAZ inhibitor K975 is capable of suppressing PCa metastasis by targeting of the CSC-like phenotypes.

**Fig. 5.**
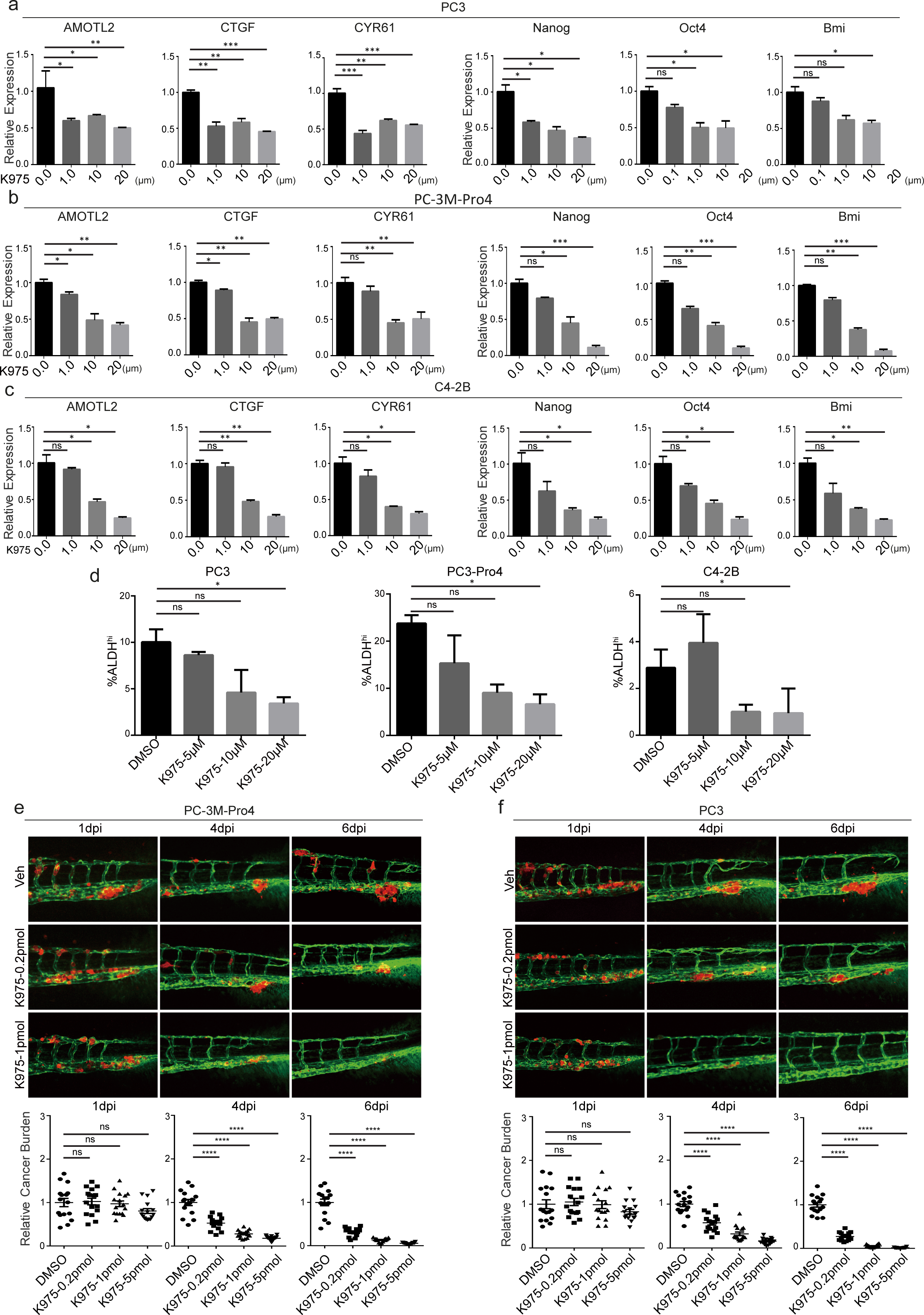
Targeting TAZ using K975 suppresses pluripotent gene expression, size of ALDH^hi^ subpopulation and metastasis in the zebrafish xenografts. (a-c) K975 treatment suppresses expression of TAZ downstream genes and pluripotent genes in PC-3, PC-3M-Pro4 and C4-2B *in vitro*. Each experiment was independently repeated 3 times. (d) K975 treatment suppresses size of the ALDH^hi^ subpopulations in PC-3, PC-3M-Pro4 and C4-2B *in vitro.* Each experiment was independently repeated 3 times. (e-f) K975 treatment suppresses metastasis of PC-3M-Pro4 (e) and PC-3 (f) in the zebrafish xenografts in a dose dependent manner. Group size=15. *p<0.05, **p<0.01, ***p<0.001, ****p<0,0001. Arrow bars are s.e.m.

### Mechanotransduction and TAZ activation promote pluripotent gene expression and organoid formation in a PCa organoid model

Having shown the significance of TAZ targeting in suppressing CSC-like phenotype and metastasis in the PCa cell lines, we next confirmed our findings using a more patient-relevant organoid model LAPC9 (48). Immunofluorescence on the organoids showed TAZ nuclear expression in cells at the peripheral cell layer (Fig. 6b), concomitant with co-localization of p-Paxillin and/or ILK with F-actin at the cell edge (Fig. 6a), implicating an involvement of the integrin-TAZ axis in formation of the organoids. We next questioned if mechanotransduction and TAZ activation are able to drive the stem-like phenotype in the LAPC9 organoids. LAPC9 cells were therefore dissociated from the organoids and seed on the elastic substrate with low (2kPa), intermediate (15kPa) and high stiffness (100kPa) for 24 hours. The cells at high stiffness showed significant expression increase of TAZ downstream genes *AMTOL2*, *CTGF* and *CYR61* and the pluripotent genes *NANOG, OCT4*, *SOX2* and *BMI1* (Fig. 6b), suggesting that organoid cells are able to sense and transduce the mechano signals (ECM stiffness) into TAZ activation and stemness gene upregulation. Unfortunately LAPC9 organoids are propagated in liquid culture and cannot be directly exposed to high stiffness therefore to further reveal the impact of TAZ activation on organoid development, we therefore overexpressed TAZ-S89A in LAPC9 cells to mimic impacts of ECM stiffness. This overexpression significantly promoted formation and growth of the organoids (Fig. 6e-f), together with increased expression of TAZ downstream genes and pluripotent genes (Fig. 6g). Overall, our studies with organoids further highlighted the importance of TAZ in stemness regulation and tumor initiation in human PCa.

**Fig. 6.**
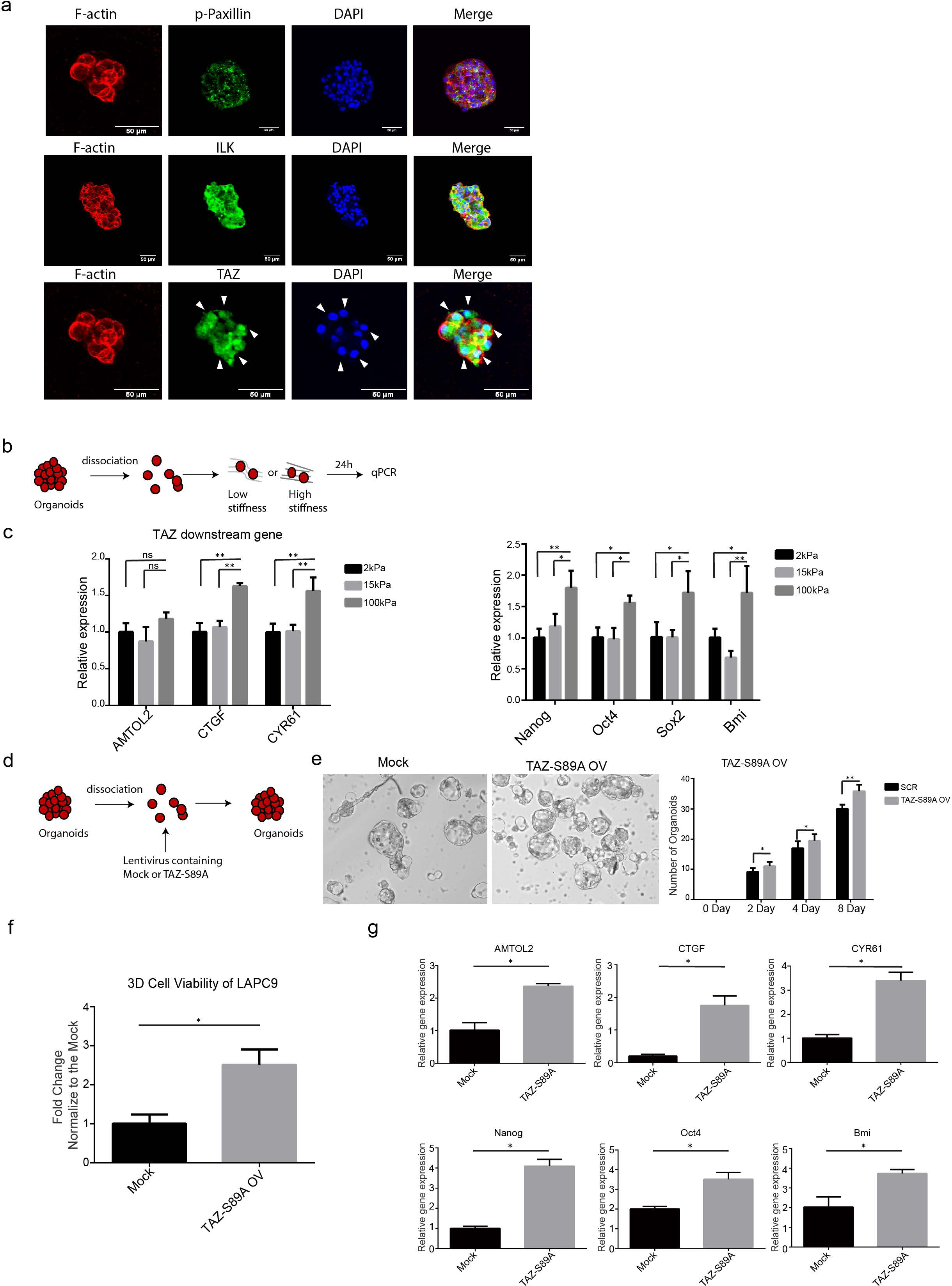
TAZ regulates pluripotent genes expression and organoid formation in LAPC9 PCa cells. (a) Subcellular localization of p-Paxillin, ILK and TAZ in LAPC9 organoids. Immunofluorescence was performed at 120 hours after seeding. Scare bar = 50um. (b-c) LAPC9 organoid cells were seeded on 2kPa, 15kPa and 100kPa for 24 hours. Expression of TAZ downstream genes and pluripotency genes were measured. Experiments were independently repeated 3 times. (d-f) Overexpression of constitutively activated TAZ increased formation of organoids (e), expression of TAZ downstream genes and pluripotent genes (f). Experiments were independently repeated 2 times. Group size = 8. *p<0.05, **p<0.01, ***p<0.001, ****p<0,0001. Arrow bars are s.e.m.

### K975 suppresses formation of PDX-derived organoids

Having shown the vital role of TAZ activation on stem gene expression and organoids formation in LAPC9 organoids, we next assessed whether pharmaceutic targeting of TAZ using K975 could reduce expression of genes indicative of cancer-stemness, and suppress formation and growth of the organoids. To achieve this, firstly, LAPC9 cells were seed in the organoid medium containing 0μM, 10 μM or 20 Μm K975 (day 0, Fig. 7a). After 72 hours of treatment, flowcytometry and qPCR were conducted to measure the expression of the CSC surface marker CD44, TAZ downstream genes *AMTOL2*, *CTGF*, *CYR61*, and pluripotent genes *NANOG* and *OCT4* (Fig. 7b-7c). Consequently, all of these markers were significantly repressed by the treatment, reflecting a reduction of cancer stemness in the organoids (Fig. 7b-7c). Next, the effect of K975 on the formation and growth of the organoids were evaluated, showing that both number and total cell titers of the organoids were inhibited (Fig. 7d-e). Overall, this data underscores the TAZ inhibitor K975 exerts strong inhibitory effect on formation and growth of the organoids.

**Fig. 7.**
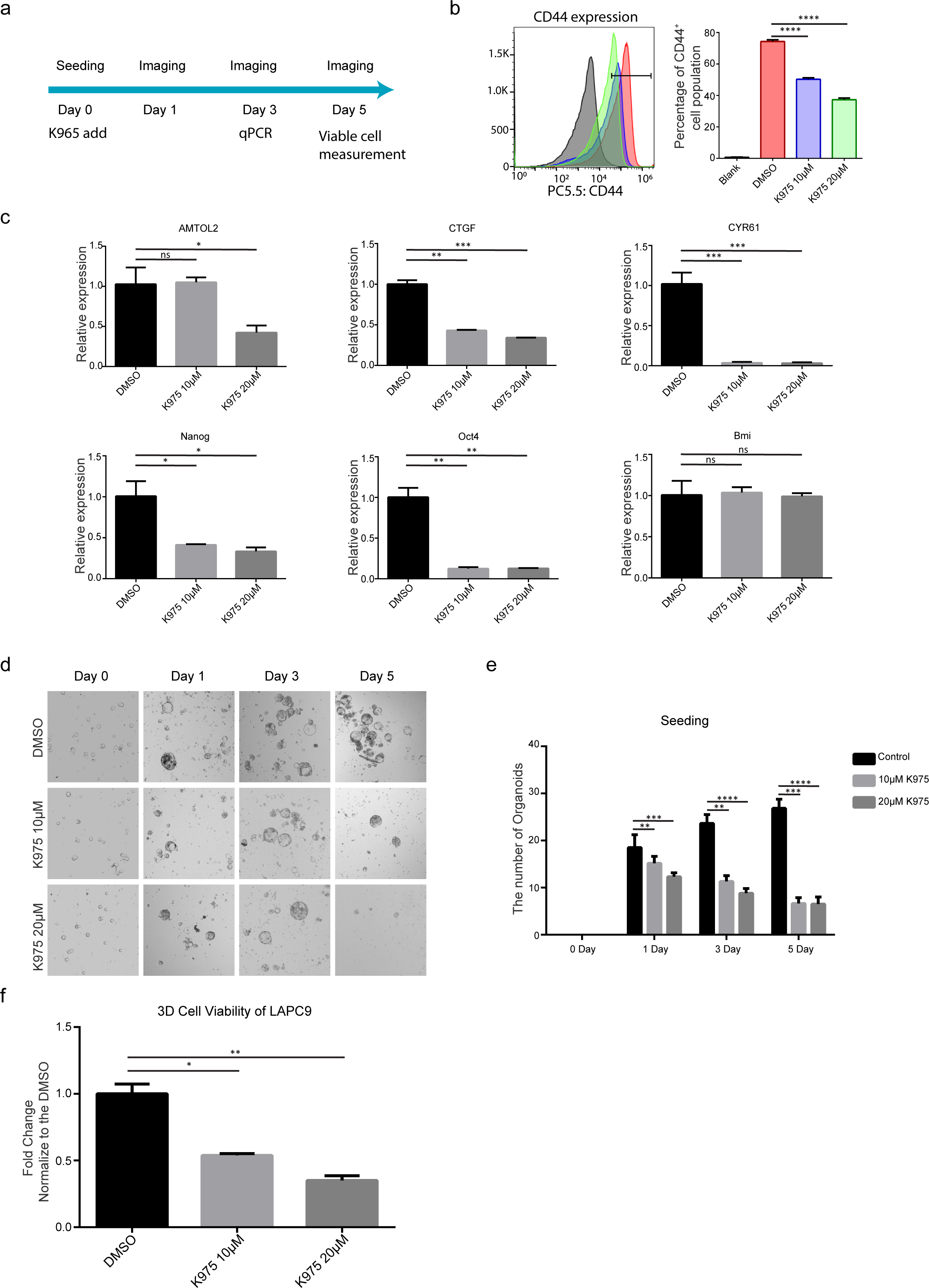
Targeting of TAZ using K975 suppresses pluripotent genes expression and organoid formation in LAPC9 cells. (a) Graphic indication for assessment of K965 on LAPC9 organoids. Single cells dissociated from LAPC9 organoids were seed in organoid medium and treated with K975. The organoids were collected at day 3 for qPCR and FACS analysis, and at day 5 for cell title analysis. (b-c) Effect of K975 on CD44 protein level (b), TAZ downstream genes expression (c, left) and pluripotent genes expression (c, right) in LAPC9 organoid after 72 hours of treatment. (d-f). Effect of K975 treatment on number and titers of LAPC9 organoids. Experiments were independently repeated 2 times. Group size = 8. *p<0.05, **p<0.01, ***p<0.001, ****p<0,0001. Arrow bars are s.e.m.

## Discussion

In prostate cancer (PCa), there is a critical need to identify druggable targets for abrogation of the heterogenic subpopulation of cells with stem-like properties that are associated with progression to lethal bone metastasis. Considering that PCa CSCs have elevated expression of cell adhesion proteins, and of the mechano-regulator TAZ (20,30,49), we investigated if stem-like PCa cells can sense and transduce matrix stiffness to facilitate their preferential homing to stiff bone. (the underlying association between mechano-transduction and CSC-like populations in PCa, and assessed anti-CSC capacity of compounds targeting the machano signaling.)

Our data indicate that the aggressive CSC-like phenotype of PCa cells relies critically on their enhanced mechanosensing and mechanotransduction potential. On one hand, the CSC-enriched ALDH^hi^ subpopulation displayed superior responses to ECM stiffness, characterized by enlarged spreading area, stronger contractile force generation and increased TAZ nuclear translocation. On the other, bulk PCa cells exposed to higher matrix stiffness exhibited significantly increased size of the ALDH ^hi^ subpopulation and enhanced pluripotent gene expression, underscoring that mechano-transduction serves as an essential driver for PCa CSC regulation. The determinant role of the mechano-signaling in CSC regulation was further revealed by gain- and loss-of-function genetic approaches in PCa cell lines as well as PDX organoids, demonstrating that the active TAZ is indispensable for maintenance of the PCa CSC-like aggressive phenotype. We further indicated that the TAZ activation in PCa is mediated by integrin-β1 and its interactor ILK, through cytoskeleton remodeling in response to mechano-stimuli (ECM stiffness). This result is in line with previous studies showing that ILK drives metastatic colonization of breast and melanoma tumor initiating cells though formation and extension of filopodia-like protrusions (FLPs) via Rif-mDIA2 and LIMK1-coffillin signaling axis (40,50). Indeed, when the ALDH ^hi^ PCa cells were cultured on matrigel, compared to their ALDH ^low^ counterparts, a significant formation of FLPs was observed accompanied by TAZ activation therefore we proposed that the Rif-mDIA2 and LIMK1-coffillin signaling axis may also engage in mechanotransduction and CSC regulation in prostate cancer.

To elucidate the importance of the mechano-signaling in PCa metastasis initiation we utilized versatile ZF xenograft model, which allows tracking of the individual cells at multiple conditions. High-resolution confocal live imaging revealed that the metastatic colonization of the injected PCa cells was accompanied by extensive actin dynamics characterized by change of cell shapes and formation of actin filaments, and TAZ activation. This actin dynamics are essential for not only cancer cell migration and extravasation but also tumor initiation and growth, as knockdowns of the actin components inhibited re-entry of cell cycle at metastatic onset (Fig. S6d). Targeting of the integrin-TAZ signaling axis also suppressed metastasis onset induced by engraftment of the ALDH^hi^ cells, suggesting this integrin-TAZ axis directly governs the aggressiveness of the CSC-like population. The critical role of the integrin-TAZ axis in PCa metastasis was further demonstrated by overexpression of ITGB1, ILK or constitutively active TAZ in PCa cells which potentiated expression of stemness genes, increased ALDH ^hi^ size and enhanced metastatic onset in zebrafish. Lastly, we tested anti-cancer efficacy of compounds putatively targeting the mechano-signaling axis, and identified the TAZ inhibitor K975 as a potential drug to suppress aggressive bone metastasis in PCa. K975 has been recently shown to have anti-cancer activity for malignant pleural mesothelioma while its efficacy on PCa remains untapped (35). In this study, we demonstrated that K975 treatment significantly suppressed expression of pluripotent genes, size of the ALDH ^hi^ subpopulations, and metastatic tumor growth in zebrafish xenografts. K975 also displayed an advanced anti-CSC activity in the PDX derived organoid model, highlighting its strong clinical potential for intervention of PCa bone metastasis, although further assessment using other pre-clinical mammalian models are required.

In conclusion, our study revealed that PCa CSCs critically respond to external stiffness and generate intrinsic forces that further enhance their pathological metastasis into stiff tissue like bone. We further presented experimental evidence that targeting of this process inhibited metastasis in zebrafish xenografts and effected development of near-patient organoids. Importantly, we selected the TAZ inhibitor K975 as a potential compound for specific targeting of PCa CSCs. Hence our study highlights the eminent role of integrin-dependent mechanotransduction in PCa metastasis initiation, which should encourage investigation of this pathway as therapeutic target in the future.

## Materials and methods

### Cell culture, *in vitro* treatment and WST assay

Human embryonic kidney cells HEK-293T (kindly provided by Dr. Sylvia Le Dévédec, LACDR, Leiden) were maintained in DMEM supplemented with 10% FCS. Human PCa cell lines PC-3 and PC-3M-Pro4-luc2 (kindly provided by Dr. Gabriel van der Pluijm, Department of Urology, LUMC) were maintained in Nutrient Mixture F-12K supplemented with 10% FCS and DMEM supplemented with 10% FCII (Hyclone™), respectively. C4-2B were maintained in low-glucose DMEM supplemented with 20% Ham’s F-12K, 10% FCS, 1x ITS Liquid Media Supplement (Thermo Fisher Scientific), 13,6pg/ml T3, 0,25ug/ml Biotin and 25ug/ml Adenine. All medium are obtained from Gibco™. For mechano-responding measurement, cells were seed on elastic cell culture substrates with defined stiffness (2, 15 or 100kPa) (ExCellness) coated with type I collagen (Sigma-Aldrich). Matrigel on top (MoT) 3D culture was performed as previously described (50). In brief, 200 cells were seed in each well of 8-well chambers coated with a thick layer of matrigel (Thermor Fisher Scientific) and covered with 200 ul full-medium containing 3% matrigel. For *in vitro* treatment, 6000 cells suspended in full medium were seed in each well of 96-well plates (6000 cells/well). After 24 hours the medium was replaced with low-serum medium (3% serum) containing Vehicle control (Veh), Integrin α2β1 inhibitor BTT3033 (TOCRIS), ILK inhibitor CPD22 (Millipore) and/or YAP/TAZ inhibitor Verteporfin (Sigma-Aldrich). Cell proliferation was assessed after 72hours of treatment using Cell Proliferation Reagent WST-1 (Roche) following the manufacturer’s protocol. 10ul WST reagent was added into each well. After 2 hours of incubation at 37 °C, the plate was measured with microplate reader M1000 PRO (TECAN). Each condition was repeated 6 times in two independent experiment.

### Cloning, lentivirus production and transduction

Short hairpin RNA (shRNA) constructs were obtained from Sigma’s MISSION library (Kindly provided by Department of Molecular Cell Biology, LUMC). pDEST/Lifeact-mCherry-N1 was a gift from Robin Shaw (Addgene plasmid # 40908), pmCherry Paxillin was a gift from Kenneth Yamada (Addgene plasmid # 50526) and pmCherry Paxillin (#50526) and pLL3.7 EGFPC2 TAZ was a gift from Yutaka Hata (Addgene plasmid # 66850). Lifeact-mCherry-N1 and mCherry-Paxillin were amplified by PCR and cloned into pLenti-blasticidin (kindly provided by Dr. Maciej Olszewski). ITGB1, ILK1 and TAZ-S89A (Addgene plasmids #32840) were amplified by PCR and cloned into pLenti-GFP-puro (Addgene plasmids #17481) Lentivirus were produced by transforming pLenti constructs, packaging plasmids psPAX2 and enveloped plasmids pMD2.G (a gift from Dr. Maciej Olszewski) into HEK-293T cells using lipoD293 (SignaGen Laboratories) as transforming reagent. Lentivirus supernatant was collected at 72hour after transformation. Cells were transduced with the lentiviruses using 6ug/ml Polybrene (Sigma-Aldrich).

### ALDEFLUOR assay and FACS sorting

The cellular subpopulation with high Aldehyde dehydrogenase (ALDH) activity was selected using ALDEFLUOR Assay kit (StemCell technology) following the manufacturer’s protocol. In brief, 1-10 million PC-3M-Pro4-Lifeact-mCherry were labelled with the ALDEFLUOR reagent. For negative control, 500,000 PC-3M-Pro4-Lifeact-mCherry cells were firstly treated with ALDEFLUOR reagent and immediately mixed with ALDH inhibitor TEAB. These cells were used to gate the negative population. FACSCanto II and FACSAria III cell sorter (BD Biosciences) were used for the measurement and sorting. Data was analyzed with FCS Express Software (De Novo Software). Each condition was independently repeated 3 times.

### Micropillar-based traction force microscopy

Cellular traction force measurements were performed using elastic micropillar arrays produced in our labs. A hexagonal array of poly-di-methyl-siloxane (PDMS, Sylgard 184, Dow Corning) micropillars of 2 um diameter, 4 um center-to-center distance and with a height of 6.1 um (Young’s modulus 28.1 kPa effective stiffness), 4.1 um (49.6 kPa effective stiffness) or 3.2 um (142 kPa effective stiffness) were produced using replica-molding from a silicon wafer (39). The pillar arrays were flanked by integrated 50 um high spacers to allow the inversion onto glass coverslips, without compromising the limited working distance of a high-NA objective on an inverted microscope. The tops of the micropillars were coated with a mixture of unlabeled and Alexa Fluor 647-labeled fibronectin (1:5, Life Technologies) using micro-contact printing. The position of the pillar tops was observed by confocal fluorescence microscopy and determined down to sub-wavelength accuracy using custom software (Matlab, Mathworks). Forces were obtained by multiplying the pillar deflections by the array’s characteristic spring constant (41.2 nN/um, 65.9 nN/um, and 191.4 nN/um, respectively, determined by finite element modeling). The pillars’ spring constants were converted to an equivalent Young’s modulus for continuous substrates (51) of 29.5 kPa (soft pillars), 47.2 kPa (medium pillars) and 137.1 kPa (stiff pillars), respectively. Only pillars closer to the cell perimeter than 3 um and with a deflection >65 nm for soft pillars, >70 nm for medium pillars, and >75 nm for stiff pillars were considered for the calculation of the total cellular forces. The deflection thresholds, which reflect the positional accuracy by which individual pillars could be localized, were determined for each confocal image as the 75th percentile of the displacements of pillars outside the cell area (i.e. not bent by the cells). The cell spreading area (A) and perimeter (p) were determined by thresholding the fluorescence signal of Alexa Fluor 532-Phalloidin labeled actin filaments using a triangular threshold method (52). The (unitless) cell elongation was calculated as *p*^2^*/(4pA)*.

Micropillar imaging was performed on a home-built setup based on an Axiovert200 microscope body (Zeiss). Confocal imaging was achieved by means of a spinning disk unit (CSU-X1, Yokogawa). The confocal image was acquired on an emCCD camera (iXon 897, Andor). IQ-software (Andor) was used for basic setup-control and data acquisition. Illumination of Alexa Fluor 532-Phalloidin and Alexa Fluor 647-fibronectin was performed with two different lasers of wavelength 514 and 642 nm (Cobolt and Spectra, respectively). Accurately controlled excitation intensity and excitation timing were achieved using an acousto-optic tunable filter (AA Optoelectronics). Light was coupled into the confocal spinning-disk unit by means of a polarization maintaining single-mode fiber (OZ Optics). The fluorescent signal was collected by a 40X/1.3 or 100X/1.4 oil objectives (Zeiss).

### Cytoimmunofluorescence

2D or 3D (MoT) cultured cells in 8 wells chamber were fixed with 4% PFA (Sigma-Aldrich) in PBS and permeablized with 0.5% triton-X 100 in PBS. After incubation with blocking buffer containing 5% BSA in PBS, the cells were incubated with primary antibody overnight at 4 °C and with fluorescent conjugated secondary antibody, Phalloidin and Dapi (Thermo Fisher) for 45 min at room temperature. Images were acquired using a Leica SP8 confocal microscope (Leica) or Nikon Eclipse Ti confocal laser-scanning microscope (Nikon). Primary antibodies used in the research included rabbit anti-human TAZ, anti-human ILK, anti-human phosphorylated Paxillin (Cell Signaling Technology) and anti-humanKi67 (Abcam). Images were processed and analyzed using imageJ software.

### Zebrafish maintenance, tumor cell implantation and metastasis analysis

Wildtype zebrafish (ZF) line ABTL and transgenic line tg (Fli:GFP) were handled in compliance with local animal welfare regulations and maintained according to standard protocols () (53).

Cancer cell transplantation was performed as described before(54–56). Briefly, two days post-fertilization (dpf), dechorionated ZF embryos were anaesthetized with 0.003% tricaine (Sigma) and plated on a Petri dish covered with 1.5% of solidified agarose. Cancer cells were trypsinized, suspended in PBS containing 2% polyvinylpyrrolidone (PVP; Sigma-Aldrich) with a concentration of 100,000 cells/ul and loaded in into borosilicate glass capillary needles (1 mm O.D. × 0.78 mm I.D.; Harvard Apparatus). 300-500 cancer cells were injected into duct of cuvier (DoC) of ZF embryos using a Pneumatic Picopump and a manipulator (WPI). The injected embryos were further maintained in a 34 °C incubator. Images were acquired with a Leica M165 FC stereo fluorescent microscope at 1-, 2-, 4- and 6-days post injection (dpi). Data was further analyzed with image J software and/or ZF4 pixel counting program (Leiden). For high resolution imaging, zebrafish embryos were placed on glass-bottom petri dishes and covered with 1% low melting agarose containing 0.003% tricaine (Sigma). Images were acquired using a Leica SP8 confocal microscope (Leica) or Nikon Eclipse Ti confocal laser-scanning microscope (Nikon) and analyzed with image J software.

### Western blot

Protein samples were collected by lysing cells with lysis buffer containing phenylmethanesulfonyl fluoride (Cell Signaling Technology) and were separated by SDS-PAGE (Bio-Rad). After transferred to polyvinylidene difluoride membranes (Millipore), the samples on the membranes were incubated with primary antibody (1:1000 times dilution) followed by horseradish peroxidase-labeled secondary antibodies (1:1000 times dilution). After treating with enhanced chemoluminescence substrate mixture (Cell Signaling Technology), blots were scanned with ChemiDoc XRS+ System (Bio-rad). Primary antibodies rabbit anti-human ITGB1, anti-human ILK, anti-human paxillin, anti-human phosphorylated YAP/TAZ (ser127/Ser89), anti-human LATS, anti-human phosphorylated LATS, anti-human MST, anti-human phosphorylated MST, anti-human Nanog and anti-human Oct-4 were obtained from Cell Science Signaling. Rabbit anti-human GAPDH and mice anti-YAP are obtained from Santa Cruz Biotechnology.

### RNA isolation and qPCR

Whole RNA was isolated using RNeasy Mini Kit (Qiagen) following the manufacturer’s introduction. cDNA synthesis and qPCR was performed using qPCR, iScript™ cDNA Synthesis Kit (Bio-rad) and iQ™ SYBR® Green Supermix (Bio-rad) following the manufacture’s protocol. Data was analyzed using 2-ΔΔct method and results were normalized to the expression level of GAPDH and/or β-actin.

### Clonogenicity assay

Single cell suspensions were seeded in 6-well plates (200 cells/well) covered with 1.5ml completed medium. After 10-14 days culturing, the cells were fixed with 4% Paraformaldehyde (PFA) and stained with 10% crystal violet (Sigma). Images were acquired using ChemiDoc XRS+ System (Bio-rad) and analyzed with image J software.

### Tumor spheroid assay

1000 single cells were suspended in 50% matrigel and seeded in a 24 wells ultra-low binding plates covered with 200uL completed medium. Number of tumor spheres was counted after 10 days culturing. experimental group.

### Organoids culture

Patient-derived PCa (CRPC) cell LAPC9 was maintained in male CB17 SCID mice. Cancer tissues were dissociated with collagenase and single cells were seeded in ultra-low attachment 6-well plates with a density of 50,000 cells/well covered with DMEM/F-12 medium supplemented with 5% FCS, Hepes, primocin, GlutaMAX, Y-27632, A83-01, SB202190, R-Spondin, Noggin, B27, N-acetyl-cysteine, Nicotinamide, EGF, F GF10, F GF2, DHT, W nt3A, HGF and PGE2. For mechanical stimulation, the organoids were dissociated using TrypLE (ThermoFisher) and seed on elastic cell culture substrates with defined stiffness (2, 15 or 100kPa) (ExCellness) coated with type I collagen (Sigma-Aldrich) and were covered with the organoid medium. After 24 hours of culture, cells were collected for qPCR.

### Statistics

Statistics analysis was performed with a Graphpad Prism 7.0 software. t-Test was used to compare two groups and ANOVA for multiple groups. Data is presented as mean ±SEM or mean ±SD. p-values ≤0.05 are considered to be statistically significant (*p≤0.05, **p<0.01, ***p<0.001, ****P<0.0001)

## Acknowledgments

We thank Dr. Gabriel van der Pluijm (Deportment of Urology, LUMC) for providing experimental material. Guido de Roo from the Flow cytometry facility (Department of Hematology, LUMC) for technical support, Martijn Rabelink and Prof. Rob Hoeben (Department of Cell Biology, LUMC) for providing lenti-viral shRNA vectors (Sigma-Aldrich). Dr. Art Jochemsen (Department of Cell Biology, LUMC) for stimulating discussion concerning TAZ inhibitor. The present work was supported by a personalized medicine grant from Alpe D’HuZes (AdH)/KWF PROPER entitled “Near-patient prostate cancer models for the assessment of disease prognosis and therapy” (UL2 014-7058). G. Zhao gratefully acknowledge the China Scholarship Council (CSC) for personal grants (No. 201906340159).

---

**Fig. S1.**
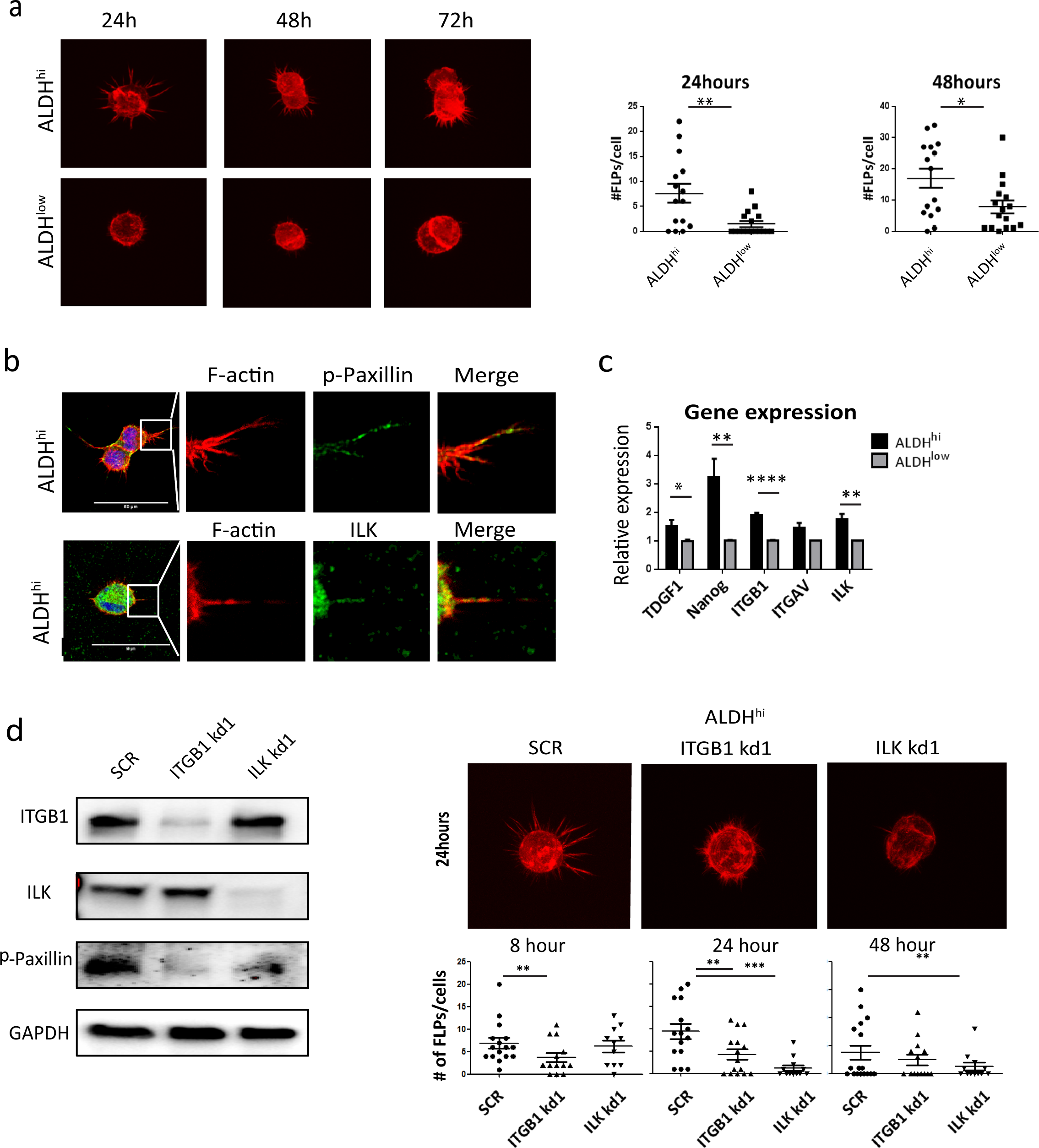
Integrin β1 and ILK regulates cytoskeleton remodeling of the ALDH ^hi^ subpopulation. (a) ALDH^hi^ subpopulation from PC-3M-Pro4 display elevated formation of filopodia-like protrusions (FLPs) than the ALDH ^low^ conterparts in Matrigel On Top (MOT) 3D culture. Experiments were independently repeated 2 times. Group size = 16. (b) Representative images for co-localization of phosphorylated Paxillin, ILK and FLPs in the ALDH ^hi^ cells in MOT 3D culture. (c) Expression of ITGB1, ITGAV, ILK, NANOG and TDGF1 in ALDH ^hi^ and ALDH ^low^ subpopulation. Experiments were independently repeated 2 times. (d) Knockdowns of ITGB1 and ILK suppress focal adhesion (marked by phosphorylation of Paxillin) and formation of FLPs in MOT 3D culture. Experiments were independently repeated 2 times. *p<0.05, **p<0.01, ***p<0.001, ****p<0,0001. Arrow bars are s.e.m.

**Fig. S2.**
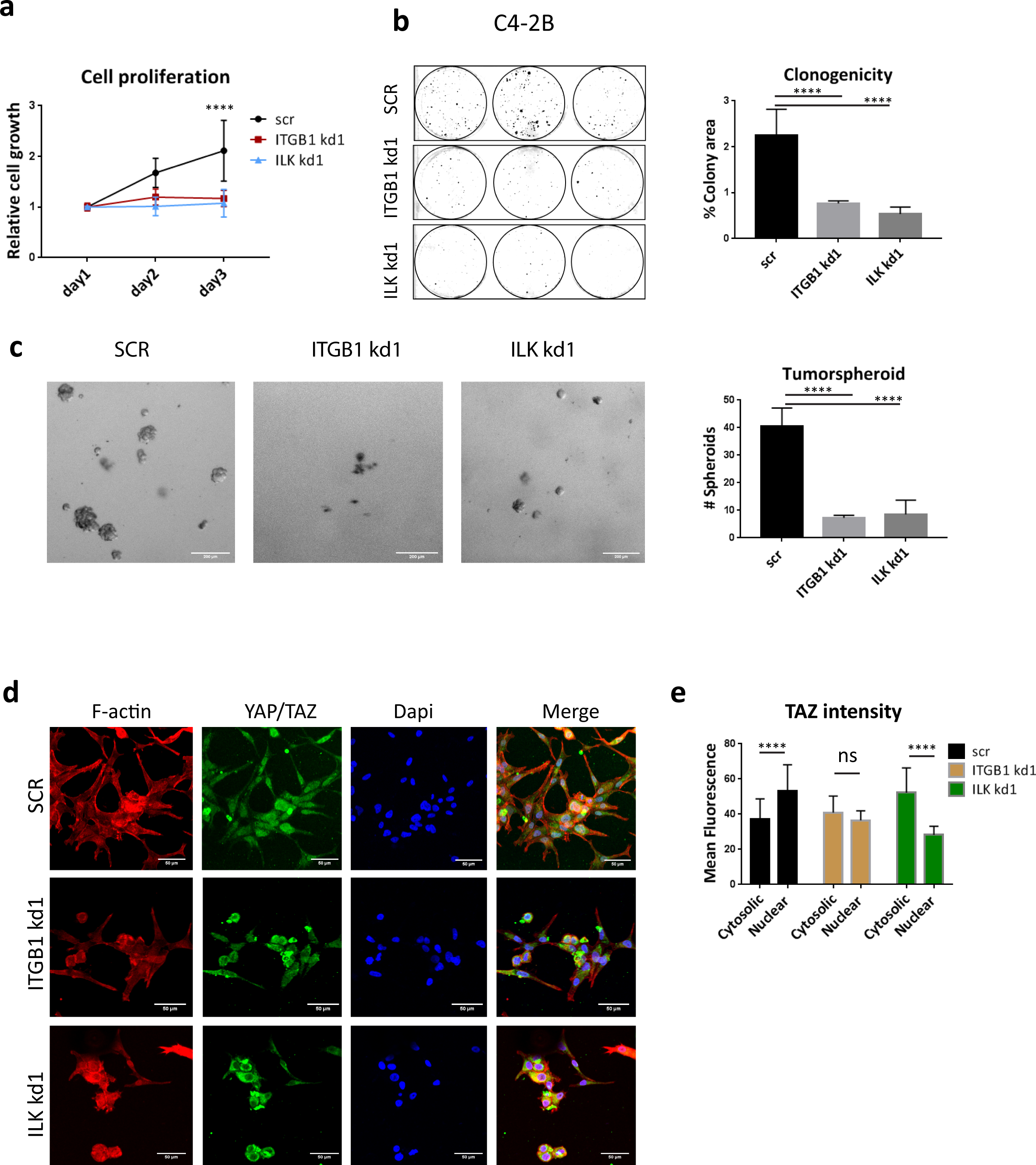
Knockdown of Integrin β1 and ILK inhibits cell proliferation, clonogenicity, tumor spheroid and YAP/TAZ nuclear translocation in C4-2B. (a) Effects of Integrin β1 and ILK knockdown on C4-2B proliferation. Experiments were repeated 2 times. (b) Effects of Integrin β1 and ILK knockdown on clonogenicity of C4-2B. 1500 cells/well were seed. Data was acquired at 14 days after seeding. Total colony area was measured using Image-J. Experiments were repeated 3 times. (c) Effects of Integrin β1 and ILK knockdown on tumor spheroid formation of C4-2B. 500 cells were suspended in 50ul of matrigel and seed in each well of 24 wells ultra-low binding plates. Numbers of tumor spheroids (>50um) were counted after 14 day culture. Scale bar = 200um. Experiment was independently repeated 3 times. (d) Effect of Integrin β1 and ILK knockdown on YAP/TAZ nuclear translocation of C4-2B on 2D culture. Cells were cultured for 48 hours. Immunofluorescence was performed against YAP/TAZ. Large bar = 50um. (e) YAP/TAZ cyotosolic/nuclear intensity was measure using Image-J. 15 cells from 2 independent experiments were analyzed.

**Fig. S3.**
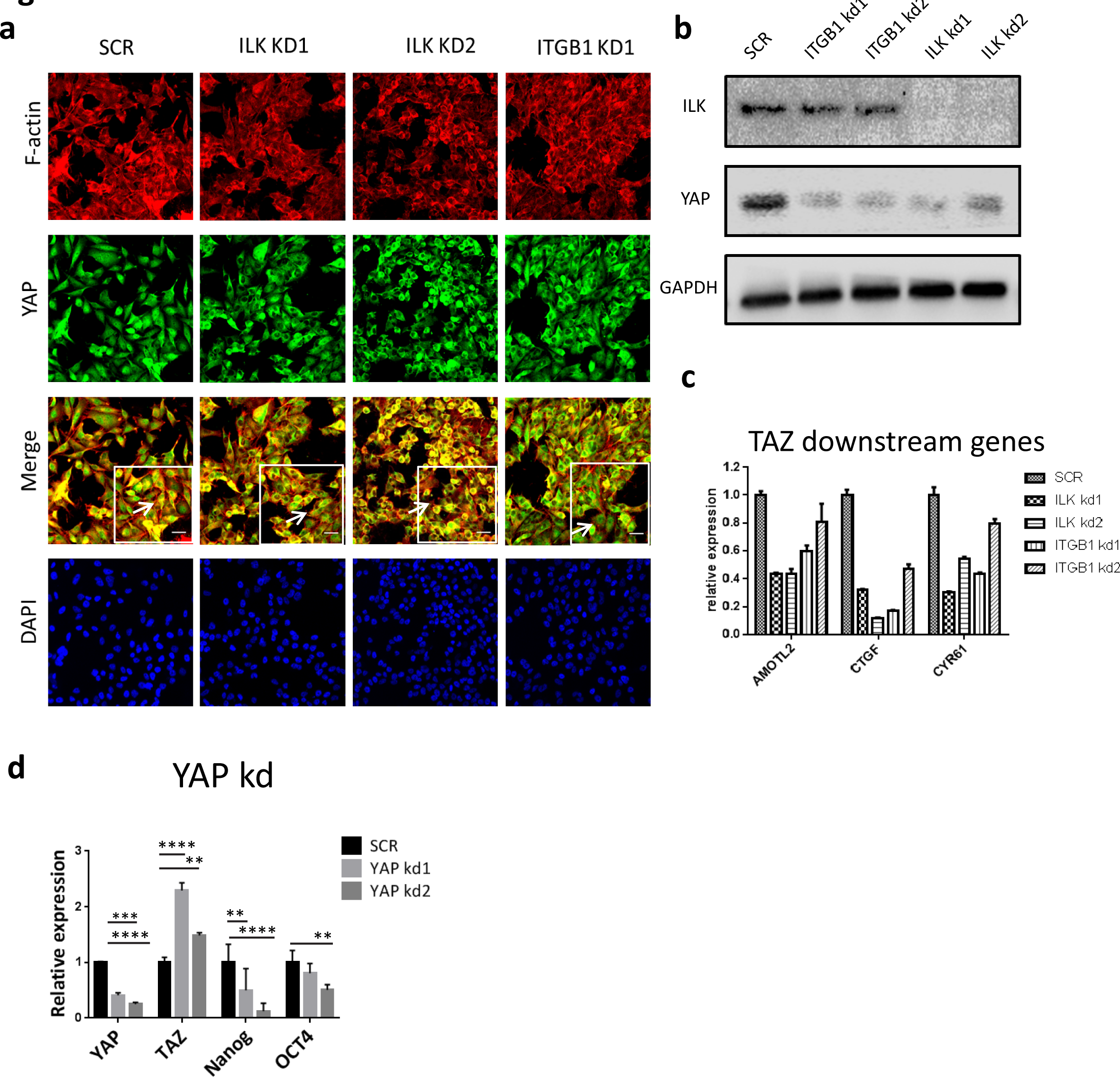
YAP is regulated by integrin β1 and ILK and is required for pluripotency gene expression and metastatic onset. (a) PC-3M-Pro4 was cultured as monolayer on glass. Immunostaining was performed against YAP. Red, F-actin; Green, YAP; Blue, Dapi. (b) Total protein level of YAP was measured in PC-3M-Pro4-SCR,-ITGB1 kd and –ILK kd using westernblot. (c) Expression of AMTOL2, CTGF and CYR61 was measured by qPCR. (d) Effects of YAP knockdowns on pluripotency gene expression.

**Fig. S4.**
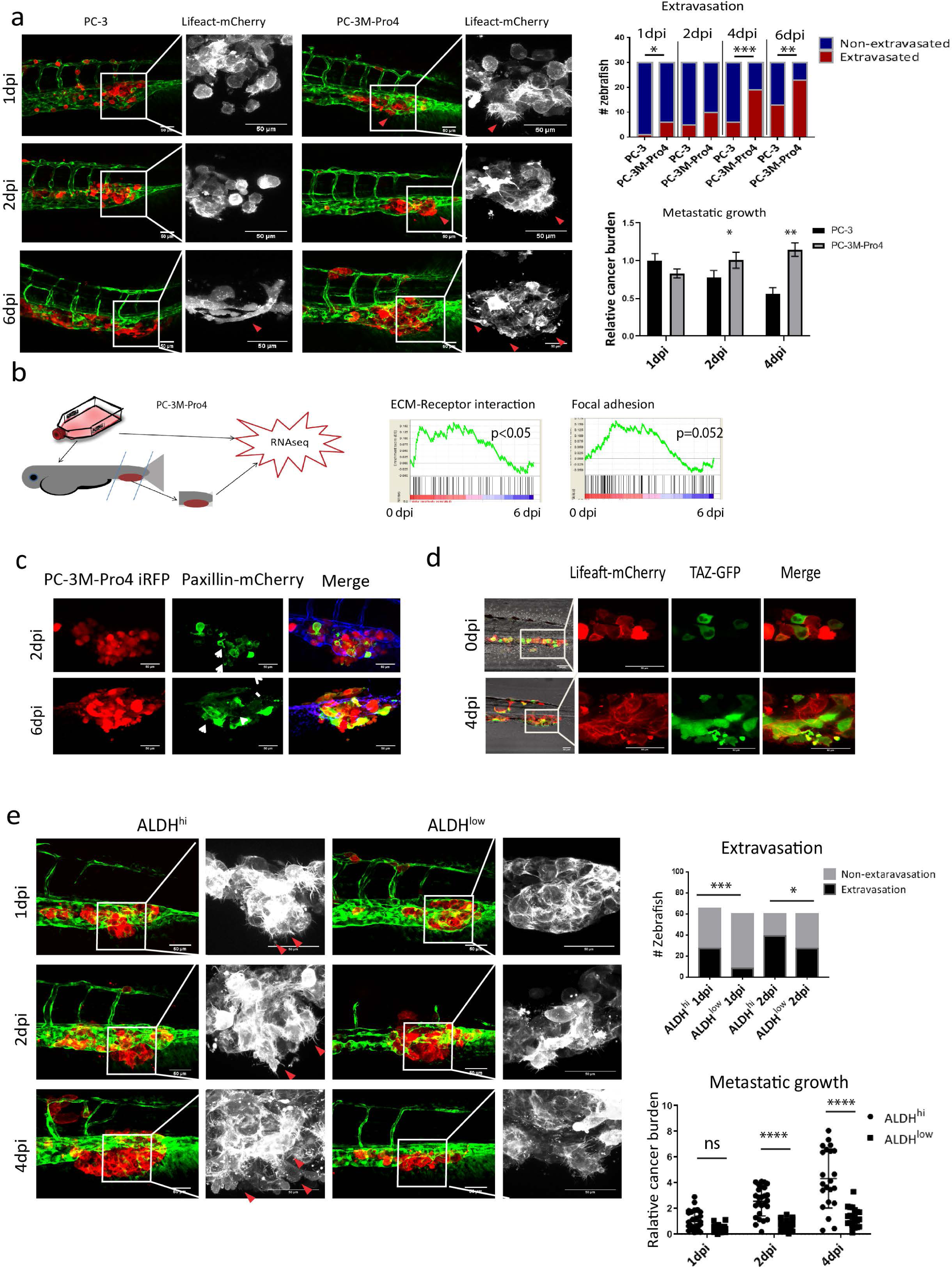
Metastatic onset of PCa cells is associated with focal adhesion, cytoskeleton remodeling and TAZ activation. (a) PC-3-Lifeact-mCherry and PC-3M-Pro4-Lifeact-mCherry were respectively injected into ZF vasculature. Metastasis at CHT was imaged using confocal at 1, 2 and 6dpi. Green, ZF vasculature; red, cancer cells; Gray, Enlargement of the cancer cells. Arrow, actin protrusions. Extravasation and total cancer cell burden were measured at 1, 2, 4 dpi. Scale bar = 50um. Group size = 30. (b) Transcriptomics of PC-3M-Pro4 was compared between in ZF metastases and in culture. Result was analyzed with Gene Set Enrichment Analysis (GSEA). (c) PC-3M-Pro4-iRFP (red) was transduced with Paxillin-mCheery (green). After injection, subcellular localization of Paxillin-mCherry was detected at 2 and 6 dpi using confocal. Scale bar = 50um. Arrow, focal adhesion plaques. (d) PC-3M-Pro4-lifeact-mCherry (red) was transduced with TAZ-GFP (green). After injection, subcellular localization of TAZ was measured using confocal. TAZ nuclear translocation was observed at 4 dpi. Scale bar = 50um. (e) ALDH ^hi^ and ALDH ^low^ cell subpopulations were isolated from PC-3M-Pro4 and injected into zebrafish. Extravasation and growth at CHT were analyzed using confocal and fluorescent microscope. ALDH^hi^ cells display enhanced extravasation and growth at the metastatic site. Group size = 30 in 2 independent experiment.. *p<0.05, **p<0.01, ***p<0.001, ****p<0,0001. Arrow bars are s.e.m.

**Fig. S5.**
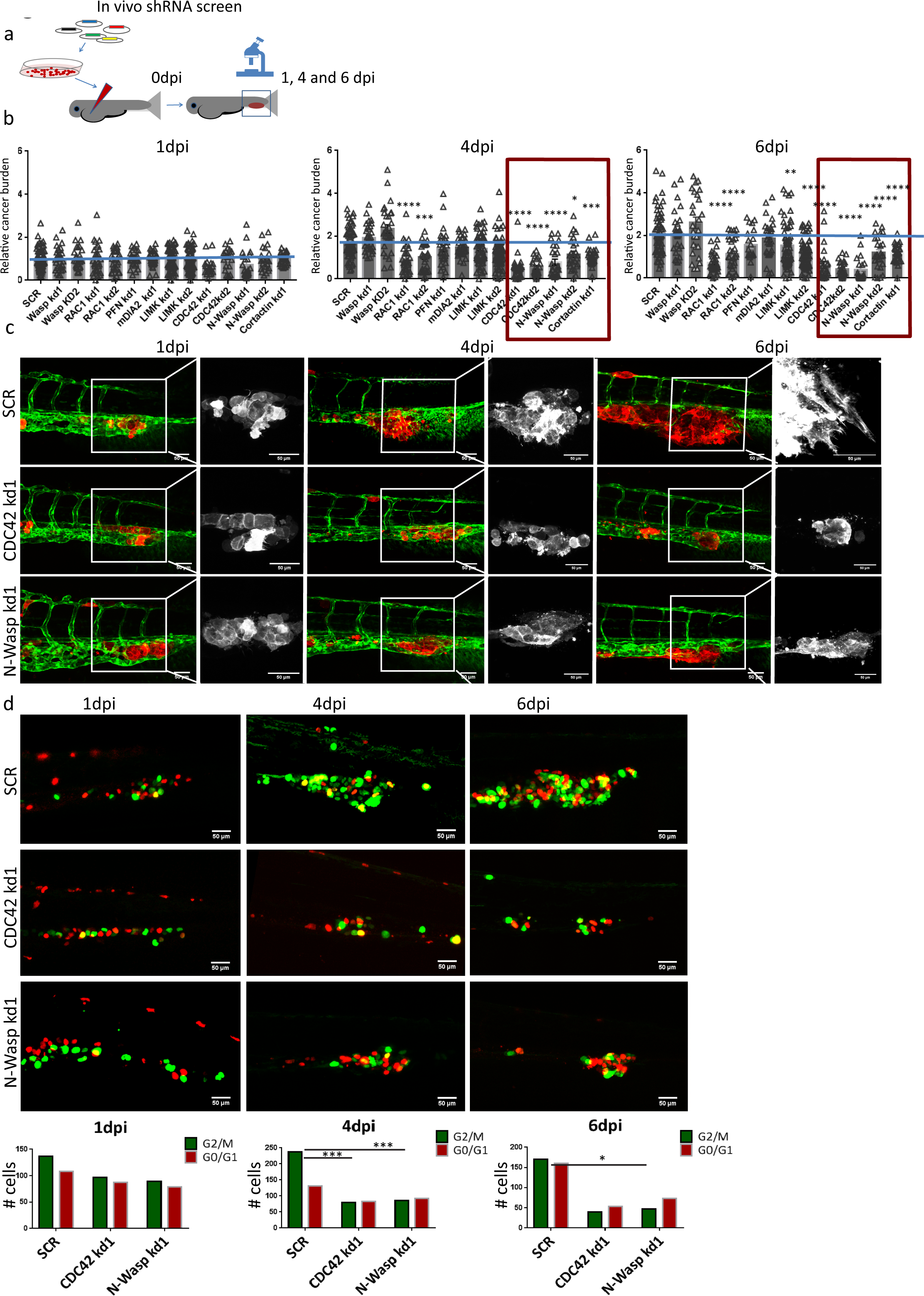
CDC42-N-Wasp-Cortactin axis is essential for metastatic outgrowth of the PCa cells. (a) Schematic indication of *In vivo* shRNA screening to identify actin factors driving PCa metastasis initiation. PC-3M-Pro4-Lifeact-mCherry containing either SCR control or shRNA against integrin-downstream actin factors were injected into zebrafish. Total cancer cell burden at metastatic site was measured at 1, 4 and 6 dpi. (b) Result of shRNA screening using zebrafish xenografts. Cancer cell burden at the metastatic site was normalized to the total fluorescent intensity at 1 dpi. Each experiment was independently repeated 2 times. Group size = 60. (c) Representative images of ZF xenografts with PC-3M-Pro4-Lifeact containing SCR, CDC42 kd1 and N-Wasp kd1 at 1, 4 and 6dpi. Scare bar = 50um. Red arrow, elongated invasive cells. (d) Effects of CDC42 and N-Wasp knockdowns on cell cycle process. PC-3M-Pro4-SCR, -CDC42kd and -N-Wasp were transduced with Fucci constructs. After transplantation into ZF, number of green and red cells were counted. 100-300 cells from 5 fish in each group were counted. Left, representative images of the engrafted cancer cells containing Fucci reporter at metastatic site, scale bar = 50um. Right, quantification. Chi-square was used for the analysis. *p<0.05, **p<0.01, ***p<0.001, ****p<0,0001. Arrow bars are s.e.m.

**Fig. S6.**
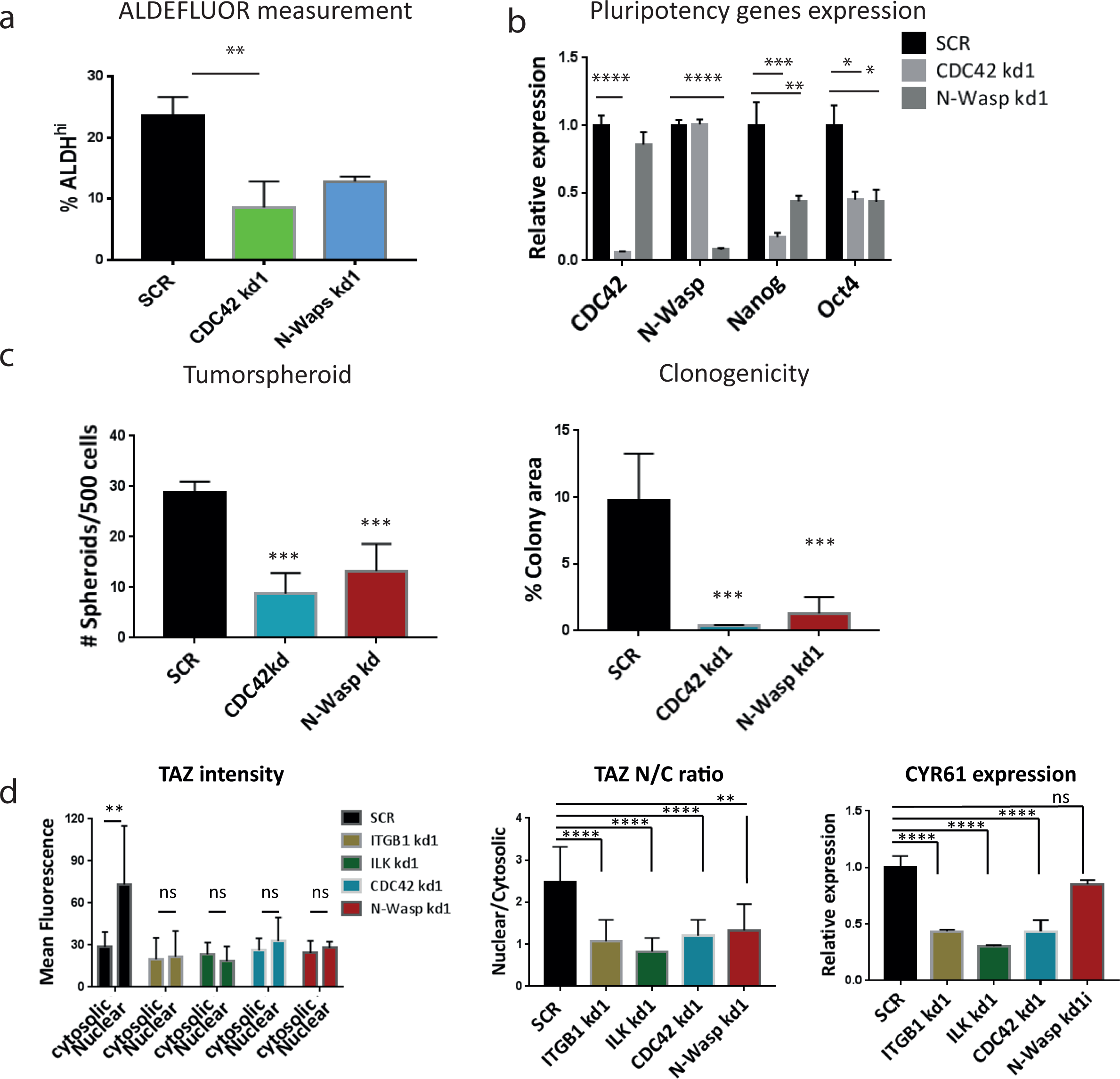
Knockdown of CDC42 and N-Wasp inhibits CSC-like properties. (a) Size of ALDH^hi^ was compared between PC-3M-Pro4-SCR, -CDC42 kd1 and -N-Wasp kd1. Experiments were independently repeated twice. (b) Expression of CDC42, N-Wasp, NANOG and OCT4 was compared between PC-3M-Pro4-SCR, -CDC42 kd1 and -N-Wasp kd1 when the cells were growing in MOT 3D culture for 96 hours. Experiments were independently repeated twice. (c) Effects of CDC42 and N-Wasp knockdowns on clonogenicity and tumorspheroid formation. 500 cells/well were seed. Data was acquired at 14 days after seeding. Total colony area was measured using Image-J. Experiments were repeated 2 times. (d) TAZ nuclear translocation and CYR61 expression were measured on the cells in MoT 3D culture. Experiments were independently repeated 2 times. *p<0.05, **p<0.01, ***p<0.001, ****p<0,0001. Arrow bars are s.e.m.

